# Impaired ability in visual-spatial attention in Chinese children with developmental dyslexia

**DOI:** 10.1101/2021.11.04.467223

**Authors:** Mengyu Tian, Runzhou Wang, Hong-Yan Bi

## Abstract

Many studies demonstrated that alphabetic language speaking children with developmental dyslexia had a deficit in visual-spatial attention, especially in rapid orienting of the attentional spotlight. Chinese, as a logographic language, is characterized as highly visual-spatial complexity. To date, few studies explored the visual-spatial attention of Chinese children with developmental dyslexia. The present study examined the visual-spatial attention of Chinese children with developmental dyslexia using the visual search task. The results showed that Chinese children with developmental dyslexia had poor performances in conjunction search, indicating that they had a deficit in the rapid orienting of visual-spatial attention. Meanwhile, only the conjunction search was a significant predictor of Chinese characters reading when other variables were controlled. These results indicated that Chinese dyslexic children had a deficit in visual-spatial attention, and visual-spatial attention played a special role in Chinese reading development.

## Introduction

Developmental dyslexia (DD) is defined as a reading skill acquisition impairment despite normal intelligence and adequate learning opportunities (Peterson & Pennington, 2012). In alphabetic languages, the phonological deficit is considered to be the most common explanation of DD (Melby-Lervåg, Lyster, & Hulme, 2012; Ramus, 2003). However, some studies found that developmental dyslexia might stem from more fundamental sensory deficits (e.g., Eden, Stein, Wood, & Wood, 1995; Tallal et al., 1996). For example, the magnocellular-dorsal pathway deficit theory is a popular model of developmental dyslexia.

The magnocellular-dorsal (MD) pathway is one of the main human visual pathways. The MD pathway begins in the retina, and carries the magnocellular information through the dorsal lateral geniculate nucleus (LGNd) to V1 and motion-sensitive visual areas (MT/V5), finally projects to the posterior parietal cortex. MD flow is thought to respond primarily to stimuli with low spatial frequencies, high temporal frequencies, and real and illusory motion (Livingstone & Hubel, 1987; Merigan & Maunsell, 1990; Schiller, Logothetis, & Charles, 1990). Although controversial opinion exists (Olulade, Napoliello, & Eden, 2013), some evidence suggests that MD deficit may be causally related to dyslexia and also be a potential cause of phonological deficits (Olulade et al., 2013). The MD pathway is also known to play a central part in visual-spatial attention, especially in directing attention to a specific location (Vidyasagar, 1999, 2005). The posterior parietal cortex (PPC) on the magnocellular–dorsal pathway is an important brain area related to visual-spatial attention (Saalmann, Pigarev, & Vidyasagar, 2007). Researchers proposed magnocellular-dorsal pathway deficits may affect reading by impairing visual-spatial attention (Vidyasagar & Pammer, 2010). When DD has a deficit in the magnocellular-dorsal pathway, the parietal lobe function is impaired due to insufficient information input from magnocellular neurons, and the visual-spatial attention ability of DD will also be impaired (Vidyasagar, 2005; Vidyasagar & Pammer, 1999, 2010).

The visual search task is often used to examine the relationship between visual-spatial attention and reading difficulty (e.g., Franceschini, Gori, Ruffino, Pedrolli, & Facoetti, 2012; Gabrieli & Norton, 2012; Liu, Chen, & Wang, 2016). There were two types of visual search tasks: parallel search and serial search task. According to the feature integration theory (FIT) (Treisman & Gelade, 1980), parallel search occurs when searching for a single feature. The search time is independent of set size (the number of items displayed). The parallel search is thought to reflect the visual processing of visual features. Simple features can be detected without attention limit, so this detection is a parallel and pre-attentive process. The conjunction search occurs when searching for two or more features in the same time. The conjunction search is a serial process that the attention spotlight focuses on a specific location at each time, and the search time increases with set size increasing. The slope of search time versus set size reflects the visual-spatial attention capacity. The serial search reflects the process of moving and focusing the “spotlight” of visual-spatial attention in space, which is related to the control of eye movements, the selection of relevant information, and the suppression of irrelevant information. This process is very similar to reading. During fluent reading, readers need visual-spatial attention spotlight sweeping from left to right within a word to segment the letter string efficiently into constituent graphemes and to choose the target letters during phonological decoding (Perry, Ziegler, & Zorzi, 2007). An fMRI study also confirmed that the activation of the parietal lobe, which is an important brain area related to visual-spatial attention increases when individuals read words letter by letter (Cohen, Dehaene, Vinckier, Jobert, & Montavont, 2008). Therefore, some researchers proposed that visual-spatial attention deficit could impede the reading acquisition through phonological decoding in alphabetic languages (Facoetti et al., 2010; Vidyasagar & Pammer, 2010). Substantial evidence showed that children with DD in alphabetic languages have deficits in visual-spatial attention, especially had difficulty in rapid orienting of the attentional spotlight (Gori, Seitz, Ronconi, Franceschini, & Facoetti, 2016; Stein, 2014; Tafti, Boyle, & Crawford, 2014; Vidyasagar & Pammer, 2010). It has been found that serial search ability is associated with rapid naming and reading speed in alphabetic languages (Di Filippo et al., 2006). Moreover, Franceschini et al. (2012) found that the serial search time in pre-reading children could predict their pseudo-word reading and length effect of pseudo-word, proving a causal relationship between visual-spatial attention and phonological decoding in alphabetic languages (Franceschini et al., 2012). Thereinto, the search time was not different between dyslexic children and normal children in the parallel search task (Sireteanu et al., 2008). But in the conjunction search task, the dyslexic children had longer search time than normal children, and their slope of the search time per item also steeper (Di Filippo et al., 2006; Sireteanu et al., 2008; Vidyasagar & Pammer, 1999; Wright, Conlon, & Dyck, 2012). These results indicated that children with dyslexia have visual-spatial attention deficits, which are not caused by visual processing difficulties.

Visual-spatial attention may be particularly important for Chinese literacy acquisition due to the special writing system (Liu et al., 2016). Unlike alphabetic languages with the linear layout of the limited number of letters, Chinese characters are the rectangular layout of radicals formed by individual strokes. Since there are a large number of similar characters in Chinese characters, more visual-spatial attention is needed to improve stimulus contrast and enhance spatial resolution to process detailed information and orthographies during Chinese character recognition, so as to accurately distinguish Chinese characters with similar shapes (Liu et al., 2016). Previous studies showed that the function of the posterior parietal cortex, which is demonstrated to be responsible for visual-spatial attention, is found to be related to orthographic skills in Chinese adults (Qian, Deng, Zhao, & Bi, 2015). The resting-state functional connectivity between the posterior parietal cortex and the left middle occipital cortex is also related to orthographic skills in Chinese adults (Qian, Bi, Wang, Zhang, & Bi, 2016). These results indicated that visual spatial attention correlated with Chinese reading skills. Moreover, the way of graphic units mapping to phonetic units is different between Chinese and alphabetic languages. Alphabetic scripts are assembled phonology, the letter by letter phonological decoding is necessary for reading. Chinese is thought to be addressed phonology because a Chinese character is directly mapping to a whole morpheme syllable (Perfetti, Cao, & Booth, 2013). A good phonological decoding skill may be less critical in Chinese reading than in alphabetic language reading (Shu, McBride-Chang, Wu, & Liu, 2006; Wei, Bi, Chen, Liu, Weng, & Wydell, 2014; Zhou et al., 2014). Above all, the cognitive demands in reading may be different between these two language systems. Indeed, Chinese children with DD are also found to show unique characteristics. The deficit in rapid naming, orthographic processing, and morphological awareness are the main problems of Chinese children with DD (Ho, Chan, Lee, Tsang, & Luan, 2004; Shu et al., 2006). Therefore, it can be speculated that Chinese children with DD may have a deficit in visual-spatial attention, and their visual-spatial attention ability should be closely related to orthographic skills rather than phonological skills. To date, only a few studies investigated the visual-spatial attention of Chinese children with DD. Ding et al. (2016) examined the visual-spatial attention in Chinese dyslexic children used two tasks: cue task and visual search task. They adopted the inhibition of return effect in eye movement as the index of visual-spatial attention. The results showed that the dyslexic children did not have the inhibition of return in the attentional task and had a visual-spatial attention deficit (Ding et al., 2016). However, in the study of Ding et al. (2016), there was no control experiment to investigate whether the observation result was caused by pure visual processing deficits. Therefore, it is still not clear whether Chinese dyslexic children have visual-spatial attention deficit after excluding the influence of visual processing deficit, and if so, how the relationship between visual-spatial attention and reading-related skills in Chinese is different from alphabetic languages.

This study aimed to examine the visual-spatial attention of Chinese dyslexic children. A serial search task, which is a conjunction search for color and orientation, was conducted to examine the visual-spatial attention in Chinese dyslexic children. We also adopted two parallel search tasks as control tasks. One was searching for color, and another was searching for orientation. If Chinese dyslexic children had visual-spatial attention deficit, they would show poorer performance in the conjunction search task. If they had difficulty in the visual processing of stimulation features, they would show poorer performance in both conjunction search and control search tasks, which is the parallel search and serial search. Finally, we also wanted to explore the relationship between visual-spatial attention and reading abilities such as rapid naming, phonological awareness, and orthographic processing skills. We hypothesized that orthographic processing skills rather than phonological processing skills in Chinese were related to visual-spatial attention.

## Method

### Participants

Forty Chinese developmental dyslexic children (10 females and 30 males, mean age is 10.99 years), and forty chronological age-matched normal children (CA, 10 females and 30 males, mean age was 11.01 years) participated in the study. All the participants were recruited from four primary schools in Beijing. They were right-handed and had normal or corrected to normal vision. They were all free of attention deficit hyperactivity disorder (based on the *Chinese Classification of Mental Disorder 3*) and neurological diseases. DD was selected based on three criteria: (1) IQ measured by *Combined Raven’s Test* was above 85. (2) The written vocabulary score measured by *Standard Character Recognition Test* (Wang & Tao, 1993) was at least 1.5 standard deviations below the average score of the same-grade children. (3) The bottom 15% of the class in reading evaluated by their teachers. CA was defined base on two criteria: (1) IQ measured by *Combined Raven’s Test* was above 85. (2) The written vocabulary measured by *Standard Character Recognition Test* was higher than the average score of the same-grade children. The two groups were matched in age, gender, and IQ, but they were significantly different in written vocabulary scores (details see Table 1). The inclusionary criteria for DD were consistent with previous studies in mainland China (Yang, Yang, Li, Xu, & Bi, 2020; Qian & Bi, 2015).

**Table 1.**
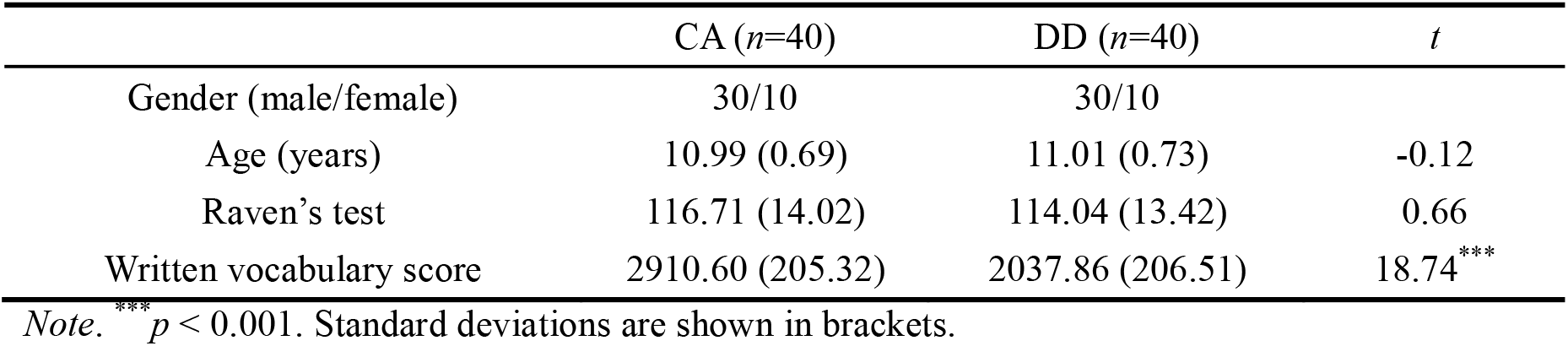
Participants’ basic information in different groups

| | CA ($n=40$) | DD ($n=40$) | $t$ |
| --- | --- | --- | --- |
| Gender (male/female) | 30/10 | 30/10 |  |
| Age (years) | 10.99 (0.69) | 11.01 (0.73) | -0.12 |
| Raven's test | 116.71 (14.02) | 114.04 (13.42) | 0.66 |
| Written vocabulary score | 2910.60 (205.32) | 2037.86 (206.51) | 18.74*** |

Note. \*\*\* $p < 0.001$ . Standard deviations are shown in brackets.

### Reading related tests

#### Reading accuracy

The same task from Qian et al. (2015) was adopted. In this task, a list of 172 characters with increasing difficulty of recognition (i.e., increasing number of strokes and decreasing word frequency) was presented on a printed paper. Participants were asked to read each character aloud in sequence without a time limit. The number of correct reading characters was recorded as the score of the test.

#### Reading fluency

The same test employed by Qian et al. (2015), which was comprised of 160 high-frequency characters. The Participants were asked to read these characters in sequence as fast as possible within one minute. The number of characters corrected read was recorded as the score of the test.

#### Rapid automatized naming test

This test included two subtests: rapid digit naming and rapid picture naming. Rapid digit naming task was from Qian et al. (2015). In this task, five digits (2, 4, 6, 7, and 9) were presented in a 6 × 5 matrix in random order. Participants were asked to name the digits as fast as possible, and each participant completed the test twice. The average time was taken as the final score. The rapid picture naming test used pictures (flower, book, dog, hand, and shoes) as materials, and it had the same process as the rapid digit naming test.

#### Phonological awareness

An oddball paradigm from Qian et al. (2015) was adopted. In this task, three Chinese single-characters were presented orally by the experimenter. The participating children were asked to point to a phonologically odd item among them. There were three types of oddity: onset, rime, and lexical tone. Ten trails for each type of oddity were presented. The number of correct responses was recorded.

#### Orthographic skill

A character judgment task from Qian and Bi (2015) was adopted. This task consisted of 40 real characters, 20 pseudo-characters, and 20 non-characters. Participants were asked to judge whether or not a presented item was a real character. The reaction time of the pseudo-characters and non-characters were referred as the index of the orthographic skills.

### Visual search

#### Stimuli

There were three types of visual search tasks in the present study, including color search, orientation search, and conjunction search for both color and orientation. Stimuli in all three visual search tasks were colored rectangles which arranged randomly in an invisible 10 × 10 grid (6.84° × 6.84° visual angle) on dark background. The rectangles were 0.3° visual angle in size. In the color search task, the target was a red rectangle, and the distractors were green rectangles. All the rectangles were horizontal. In the orientation search task, the target was a horizontal rectangle, and the distractors were vertical rectangles. All rectangles were green. In the conjunction search task, the target was a red vertical rectangle, and the distractors were green vertical rectangles and red horizontal rectangles.

#### Procedure

Participants performed the three visual search tasks in a random order, and there were four blocks with varied set sizes of 12, 24, 36 and 48 in each task. In color and orientation search tasks, each block contained 12 trials with the same set size. In the conjunction search task, each block contained 24 trials with the same set size. Half of the trials in one block contained the target and the other trails did not have. Four blocks in each task were presented randomly. Each trial was started with 500 ms fixation, and then the stimulus was presented until the participant responded. Participants were asked to judge whether the target was present or not and press the keys (“f” or “j”) to respond. The responding keys of target-present and target-absent were balanced across participants. The participants were instructed to respond as quickly as possible without sacrificing accuracy. Accuracy and reaction times (RTs) were recorded.

## Results

### Performances of reading skills

The results of reading skills are shown in Table 2. Children with dyslexia had poor performances than typically developing children in all tests. Specifically, the DD group had longer reaction times on no-characters than CA group, but there was no difference between DD and CA group in reaction times on pseudo-characters.

**Table 2.** Performance in all tests for children with and without dyslexia

|  | CA (*n*=40) | DD (*n*=40) | *t* |
| --- | --- | --- | --- |
| Reading accuracy (items) | 105.5 (10.07) | 83.17 (11.99) | 8.83<sup>***</sup> |
| Reading fluency (items) | 104.97 (18.13) | 72.61 (15.76) | 8.36<sup>***</sup> |
| Digit rapid naming (s) | 9.3 (2.27) | 12.27 (2.53) | -3.84<sup>***</sup> |
| Picture rapid naming (s) | 17.43 (4.28) | 20.97 (3.74) | -5.31<sup>***</sup> |
| Phonological awareness (items) | 24.21 (4.26) | 18.82 (6.03) | 4.51<sup>***</sup> |
| RT of pseudo characters (ms) | 952.10 (230.02) | 1015.15 (374.74) | -0.87 |
| RT of false characters (ms) | 767.52 (137.61) | 831.01 (126.70) | -2.07<sup>**</sup> |

*Note.* \* $p < 0.05$ , \*\*\* $p < 0.001$ . Standard deviations are shown in brackets.

### Visual search

The participants whose accuracy was lower than 50% were excluded, and 39 CA and 36 DD in color search task, 39 CA and 38 DD in orientation search task, and 34 CA and 34 DD in conjunction search task were remained. The trials with wrong reactions or with RTs longer than three standard deviations away from the mean (less than 1%) were also excluded. The accuracy and the RTs of the three tasks were respectively shown in Table 3 and Table 4.

**Table 3.** The accuracy in three visual search tasks

|  | Set size | DD |  | CA |  |
| --- | --- | --- | --- | --- | --- |
|  |  | With target | Without target | With target | Without target |
| Color search | 12 items | 0.91 (0.10) | 0.96 (0.09) | 0.95 (0.08) | 0.95 (0.08) |
|  | 24 items | 0.97 (0.06) | 0.95 (0.09) | 0.95 (0.09) | 0.96 (0.07) |
|  | 36 items | 0.92 (0.08) | 0.97 (0.07) | 0.96 (0.10) | 0.98 (0.06) |
|  | 72 items | 0.95 (0.08) | 0.98 (0.05) | 0.95 (0.11) | 0.97 (0.06) |
| Orientation search | 12 items | 0.92 (0.10) | 0.95 (0.09) | 0.96 (0.07) | 0.95 (0.08) |
|  | 24 items | 0.96 (0.08) | 0.95 (0.10) | 0.97 (0.06) | 0.97 (0.07) |
|  | 36 items | 0.96 (0.08) | 0.96 (0.08) | 0.95 (0.08) | 0.97 (0.08) |
|  | 72 items | 0.96 (0.08) | 0.98 (0.05) | 0.97 (0.07) | 0.97 (0.06) |
| Conjunction search | 12 items | 0.87 (0.10) | 0.97 (0.07) | 0.91 (0.09) | 0.98 (0.04) |
|  | 24 items | 0.86 (0.14) | 0.97 (0.04) | 0.89 (0.13) | 0.97 (0.06) |
|  | 36 items | 0.79 (0.17) | 0.99 (0.02) | 0.84 (0.14) | 0.98 (0.04) |
|  | 72 items | 0.70 (0.16) | 0.95 (0.13) | 0.77 (0.16) | 0.99 (0.02) |

*Note.* The accuracy reported in the percentage of correct trials. Standard deviations are shown in brackets.

**Table 4.**
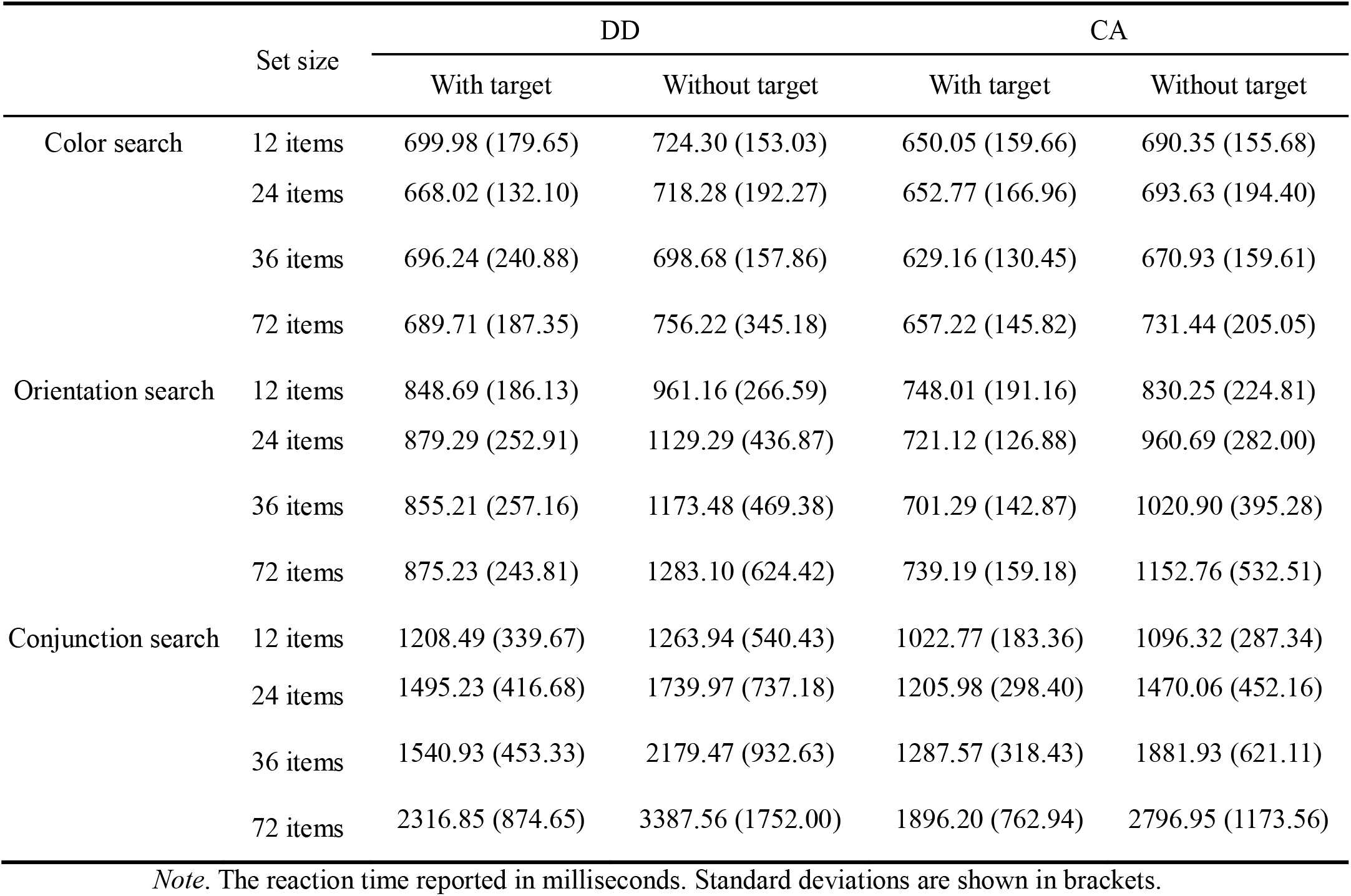
The RTs in three visual search tasks

Three-way mixed ANOVAs were conducted on accuracy in three visual search tasks separately, with group (CA, DD) as a between-subject factor and target (present, absent) and set size (12, 24, 36, 72 items) as two within-subject factors. Greenhouse-Geisser correction was reported whenever Mauchly’s test of sphericity was significant. In the color search task, the main effect of target was significant [*F* (1, 73) = 5.41, *p* < 0.05, η_p_^2^ = 0.07], accuracy was significantly higher in target-absent trials than that in target-present trials. Other main effects and any interactions were not significant, all *F*s < 1, all *p*s > 0.05. In the orientation search task, only the main effect of set size was significant [*F* (3, 225) = 2.80, *p* < 0.05, η_p_^2^ = 0.04]. Post hoc analysis with Bonferroni correction for multiple comparisons showed the accuracy was lower in the 12 items trials than in the 72 items trials. There was no significant difference of accuracy between any two other pairs of set sizes. There was no other main effect and no significant interactions (*p*s > 0.05). In the conjunction search task, the main effect of target was significant [*F* (1, 66) =136.96, *p* < 0.001, η_p_^2^ = 0.68], which accuracy was significantly higher in target-absent trials than that in target-present trials. The main effect of set size was significant [*F* (3, 198) = 25.36, *p* < 0.001, η_p_^2^ = 0.26], which the accuracy decreased with the set size increasing (12 items vs 36 items: *p* = 0. 010; 12 items vs 72 items: *p* < 0.001; 24 items vs 72 items: *p* < 0.001; 36 items vs 72 items: *p* < 0.001), but no significant difference between 12 items and 24 items (*p* = 1.000), and between 24 items and 36 items (*p* = 0.220). The main effect of group was significant [*F* (1, 66) = 5.40, *p* < 0.05, η_p_^2^ = 0.08], which children in CA group have higher accuracy than children in DD group. The interaction between target and set size was significant [*F* (2.37, 156.43) = 16.55, *p* < 0.001, η_p_^2^ = 0.20]. The results of simple effect analysis showed that in target-present trials, the accuracy decreased with the set size increasing (12 items vs 36 items: *p* = 0. 001; 12 items vs 72 items: *p* < 0.001; 24 items vs 36 items: *p* = 0.035; 24 items vs 72 items: *p* < 0.001; 36 items vs 72 items: *p* = 0.005), but no significant difference between 12 items and 24 items (*p* = 1.000). However, in target-absent trials, the accuracy was not significantly different among the four set-size conditions.

The accuracy of the target-absent condition was significantly higher than the target-present condition in color search and conjunction search. Meanwhile, the RT of the target-absent condition was significantly slower than the target-present condition in color search [710.48 ms vs 667.90 ms, *F* (1, 73) = 13.99, *p* < 0.001, η_p_^2^ = 0.16] and conjunction search [1977.03 ms vs 1496.76 ms, *F* (1, 66) = 80.81, *p* < 0.001, η_p_^2^ = 0.55]. These results demonstrated that the participants may have sacrificed fast responses for higher accuracy, especially in the target-absent condition, as a manifestation of the speed-accuracy trade-off. Thus, scaled RTs (Madden et al., 2017) were calculated as the participant’s RTs divided by the accuracy of that trial type and used in the further analysis (details see Table 5).

**Table 5.** The scaled RTs in three visual search tasks

|  | Set size | DD |  | CA |  |
| --- | --- | --- | --- | --- | --- |
|  |  | With target | Without target | With target | Without target |
| Color search | 12 items | 749.30 (225.97) | 764.61 (197.01) | 691.13 (240.64) | 726.93 (180.64) |
|  | 24 items | 706.66 (163.04) | 733.78 (199.72) | 704.74 (270.01) | 726.17 (219.75) |
|  | 36 items | 735.75 (252.42) | 732.10 (199.84) | 656.18 (141.67) | 685.68 (174.42) |
|  | 72 items | 733.26 (211.97) | 777.86 (353.20) | 706.62 (218.74) | 754.73 (206.88) |
| Orientation search | 12 items | 927.60 (251.15) | 1007.85 (267.73) | 780.46 (202.27) | 866.95 (213.88) |
|  | 24 items | 914.96 (257.93) | 1186.33 (425.06) | 745.75 (136.69) | 983.75 (271.10) |
|  | 36 items | 894.38 (301.85) | 1225.88 (521.34) | 733.93 (144.22) | 1050.87 (390.42) |
|  | 72 items | 927.90 (346.92) | 1245.87 (594.40) | 755.47 (152.24) | 1189.44 (553.04) |
| Conjunction search | 12 items | 1387.03 (414.11) | 1316.96 (619.74) | 1132.11 (256.62) | 1112.57 (288.91) |
|  | 24 items | 1767.06 (566.10) | 1782.68 (738.77) | 1407.93 (532.64) | 1502.43 (448.17) |
|  | 36 items | 2011.99 (731.63) | 2191.69 (925.62) | 1594.09 (643.90) | 1922.59 (684.66) |
|  | 72 items | 3443.99 (1725.89) | 3554.25 (1713.87) | 2606.05 (1337.38) | 2803.08 (1164.94) |

*Note.* The reaction time reported in milliseconds. Standard deviations are shown in brackets.

The same ANOVAs were conducted for the scaled RTs in three visual search tasks separately. Greenhouse–Geisser correction was reported whenever Mauchly’s test of sphericity was significant.

In the color task, the main effect of target was significant [*F* (1, 73) = 4.38, *p* < 0.05, η_p_^2^ = 0.06]. Scaled RTs in target absent condition were longer than that in target present condition. Neither the main effect of set size [*F* (3, 219) = 1.48, *p* = 0.221] nor the main effect of group [*F* (1, 73) = 0.86, *p* = 0.358] was significant. There were also no significant interactions between group and target [*F* (1, 73) = 0.24, *p* = 0.623], between group and set size [*F* (3, 219) = 0.77, *p* = 0.514], and among group, set size and target [*F* (2.54, 185.18) = 0.16, *p* = 0.895].

In the orientation task, the main effect of group was significant [*F* (1, 75) = 9.64, *p* < 0.01, η_p_^2^ = 0.11], children in DD group took a longer time to respond than children in CA group. The main effect of target was significant [*F* (1, 75) = 53.52, *p* < 0.001, η_p_^2^ = 0.42], scaled RTs in target-absent condition were longer than that in target-present condition. The main effect of set size was significant [*F* (2.48, 185.95) = 8.47, *p* < 0.001, η _p_^2^ = 0.10], scaled RTs in condition of the set size with 12 and 24 items were significant shorter than that with 72 items (*ps* < 0.05). The interaction between target and set size was significant [*F* (2.40, 179.69) = 14.11, *p* < 0.001, η_p_^2^ = 0.16]. The results of simple effect analysis showed that there is no significant difference among any two scaled RTs of four types of set size in target present condition (*p*s > 0.05),but in target absent condition, the scaled RTs in condition of the set size with 12 items were significant shorter than that with other items (*ps* < 0.001), and scaled RTs in condition of the set size with 24 items were significant shorter than that with 72 items (*p* < 0.05). There were no other interactions.

In the conjunction task, the main effect of group was significant [*F* (1, 66) = 6.71, *p* < 0.05, η_p_^2^ = 0.09], the scaled RTs of DD group were longer than CA group. The main effect of set size was significant [*F* (1.35, 88.94) = 154.76, *p* < 0.001, η_p_^2^ = 0.70], the scaled RTs increased with the set size. Specifically, the scaled RTs of 12 items were shorter than 24 items (*p* < 0.001), 36 items (*p* < 0.001), and 72 items (*p* < 0.001); the scaled RTs of 24 items were shorter than 36 items (*p* < 0.001), and 72 items (*p* < 0.001); the scaled RTs of 36 items were shorter than 72 items (*p* < 0.001). The interaction effect between set size and group was significant [*F* (1.35, 88.94) = 3.83, *p* < 0.05, η_p_^2^ = 0.06]. Simple effect analysis with Bonferroni correction for multiple comparison showed that the scaled RTs of CA group were shorter than that of DD groups in 12 items trials (mean difference = −229.66 ms, *p* = 0.017), 24 items trials (mean difference = −319.69 ms, *p* = 0.015), 36 items trials (mean difference = −343.50 ms, *p* = 0.026) and 72 items trials (mean difference = −794.55 ms, *p* = 0.019). The mean difference between two groups increased with the set size increasing.

In order to measure the overt attention more clearly, this study referred previous studies (e.g., Casco & Prunetti, 1996; Li, Sampson, & Vidyasagar, 2007) to calculate the slope of the regression line of the scaled RTs functions over set size in conjunction search for each participant. The slope was analyzed in a two-way mixed ANOVA with group (CA, DD) as a between-subject factor, and target (present, absent) as a within-subject factor. The main effect of group was significant [*F* (1, 66) = 4.42, *p* < 0.05, η_p_^2^ = 0.06], the slope of DD group was deeper than CA group. The main effect of the target [*F* (1, 66) = 1.18, *p* = 0.282] and the interaction between the group and target were not significant [*F* (1, 66) =0.01, *p* = 0.930].

## The relationship between reading skills and the ability of visual search

Partial correlations across all tasks, with gender, nonverbal IQ, and age statistically controlled, are presented in Table 6. The reaction time of the orthographic skills (Orthographic RT) was the reaction time of the pseudo-characters minus the non-characters. Fifty-six participants were included in correlation analysis because they completed all the tests. In order to reflect the ability of visual search, the slope of the regression line of the scaled RTs (in the target-present condition) functions over set size in three search tasks were calculated for each participant. Rapid picture naming was significantly correlated with color search ability. Notably, the reaction time of the orthographic skills was significantly correlated with conjunction search ability.

**Table 6.** Correlations between the ability in search tasks and reading-related scores

| Variable | 1 | 2 | 3 | 4 | 5 | 6 | 7 | 8 | 9 |
| --- | --- | --- | --- | --- | --- | --- | --- | --- | --- |
| 1. Reading fluency | – |  |  |  |  |  |  |  |  |
| 2. Reading accuracy | 0.79** | – |  |  |  |  |  |  |  |
| 3. Phonological awareness | 0.31* | 0.42** | – |  |  |  |  |  |  |
| 4. Rapid picture naming | -0.54** | -0.48** | -0.30* | – |  |  |  |  |  |
| 5. Rapid digit naming | -0.70** | -0.62** | -0.44** | 0.64** | – |  |  |  |  |
| 6. Orthographic RT | -0.07 | -0.04 | 0.05 | 0.16 | 0.09 | – |  |  |  |
| 7. Slope (color search) | 0.07 | 0.03 | 0.07 | -0.28* | -0.22 | 0.08 | – |  |  |
| 8. Slope (orientation search) | -0.03 | -0.04 | -0.16 | 0.12 | 0.05 | -0.15 | 0.03 | – |  |
| 9. Slope (conjunction search) | -0.11 | -0.23 | 0.14 | 0.03 | 0.09 | -0.34* | 0.24 | 0.05 | – |

\* $p < 0.05$ , \*\* $p < 0.01$ , \*\*\* $p < 0.001$

Hierarchical regression analyses were used to examine the relationships between reading fluency, reading accuracy and visual searching abilities after controlling for gender, nonverbal IQ, age, and reading-related cognitive abilities. Gender, age, and IQ were first entered as covariates in Step 1. Then, reading-related cognitive abilities were entered in Step 2. Finally, the slopes of three search tasks were stepwise regression in Step 3. The results of the hierarchical regression analyses indicated that only the conjunction search was a significant predictor of Chinese word reading when other variables were controlled. In particular, the conjunction search accounted for 5% of the significant variance in reading accuracy (see Table 7).

**Table 7.** Hierarchical Regression of three search tasks on reading abilities

| Step | | Cumulative $R^2$ | $R^2$ change |
| --- | --- | --- | --- |
| *Reading fluency* |  |  |  |
| 1 | Age, Gender, IQ | 0.02 | 0.02 |
| 2 | Reading-related skills | 0.51 | 0.49<sup>***</sup> |
| *Reading accuracy* |  |  |  |
| 1 | Age, Gender, IQ | 0.04 | 0.04 |
| 2 | Reading-related skills | 0.44 | 0.40<sup>***</sup> |
| 3 | Conjunction search | 0.49 | 0.05<sup>*</sup> |

\* $p < 0.05$ , \*\* $p < 0.01$ , \*\*\* $p < 0.001$

## Discussion

This study using visual search task examined the visual-spatial attention, especially the rapid orienting of the attentional spotlight in Chinese dyslexic children. The results showed that Chinese dyslexic children had poorer performance in both orientation search and conjunction search tasks, except color search tasks. The results of correlate analysis showed that the performance in the color search task was correlated with the rapid picture naming, and the conjunction search performance was correlated with the Orthographic RT. Finally, only the conjunction search ability was a significant predictor of Chinese word reading when other reading-related factors were controlled.

### Pure visual processing abilities in Chinese dyslexia

The result showed that the performances of Chinese dyslexic children were poorer than the control group in orientation search, suggesting that Chinese dyslexic children might have a deficit in the visual processing of the orientation, at least in vertical and horizontal. In alphabetic languages, researchers found that dyslexic children did not show a deficit in orientation search or shape search (Sireteanu et al., 2008; Vidyasagar & Pammer, 1999). The Chinese character may have more visual processing demand than alphabetic language. A fMRI study found that reading Chinese had stronger activations in the early visual cortices than reading alphabetic language (Szwed, Qiao, Jobert, Dehaene, & Cohen, 2014). These results suggested that the deficit in the visual processing of the orientation might be specific to Chinese children with DD. Chinese characters are formed by strokes. In many cases, the difference between the two characters is subtle. For example, there is only a slight difference between the two characters “睛”, which means the “eye”, and “晴”, which means “sunshine”. Moreover, both characters included several horizontal and vertical strokes. Thus, fast and accurate visual processing of horizontal and vertical lines (e.g., orientation search) plays an essential role in the development of Chinese reading. Furthermore, some researchers proposed that bottom-up visual attention also involved in parallel search (Wolfe, 1994). Lin, Wang, and Meng (2013) found that there was no difference between dyslexia and control group in the simple visual search task (color search). However, children with dyslexia had significantly longer reaction time than the control group to retrieve the target stimuli in complex visual search task (ellipse search). Lin and his colleagues suggested that an important difference between complex visual search and the simple visual search was that complex visual search required meticulous processing of target and interference through attention. In the present study, the stimuli used in the orientation search were similar to those used in the elliptic search. Therefore, the poor performances of Chinese dyslexic children in orientation search could be attributed to a deficit in the bottom-up visual attention.

### Rapid orienting of the visual-spatial attention spotlight in Chinese dyslexia

The conjunction search is a serial process that needs the visual-spatial attention spotlight (Quinlan, 2003; Treisman & Gelade, 1980). We found the Chinese children with DD showed poor performance in conjunction search, indicating that Chinese children with DD have a deficit in the rapid orienting of the visual-spatial attention spotlight. It is similar to the findings in alphabetic language speaking children with DD (Di Filippo et al., 2006; Sireteanu et al., 2008; Vidyasagar & Pammer, 1999; Wright et al., 2012). Furthermore, the present study found that the conjunction search ability was a significant predictor of Chinese word reading when other variables were excluded. The performance in conjunction search was correlated with the RT of orthographic skill, and no other correlation was found, including phonological awareness. These results suggested the way that visual-spatial attention influenced the reading skills might be different between Chinese and alphabetic language because the way of graphic units mapped to phonetic units in Chinese is different from alphabetic language. In alphabetic languages, evidence was found that the visual-spatial deficit was correlated with the phonological decoding deficit (Facoetti et al., 2006; Facoetti et al., 2010; Franceschini et al., 2012; Franceschini et al., 2013; Franceschini et al., 2017). The alphabetic language is the assembled-phonology, but the Chinese are the addressed-phonology. There is no letter by letter decoding when Children learn to read Chinese characters. Therefore, visual-orthographic skill is important in visual-spatial attention due to the high visual complexity of Chinese characters (Liu et al., 2016; Liu, Liu, Pan, & Xu, 2018). There are a large number of Chinese characters that differ only by one stroke or a couple of strokes, e.g., 王/wang2/ (king) and 工/gong1/ (work). In order to process visually similar characters, children must direct their visual-spatial attention to recognize subtle stroke differences (Liu et al., 2016). Unlike the alphabetic language, a Chinese character is an independent visual unit, and there is no interword spacing for multicharacter words. Thus, adequate visual-spatial attention may help the reader identify words in a sentence with ambiguous word boundaries (Liu, Chen, & Chung, 2015). For example, in the sentence “ 过几天气温会逐渐升高” (the temperature will gradually increase in a few days), “ 气” /qi4/ forms the word “气温” /qi4 wen1/ (temperature) with the character that comes after it. Although “气” can also combine with the character that comes before it to form “天气” /tian1 qi4/ (weather), in this particular sentence context, “气温” is a word, whereas “天气” is not. Therefore, visual-spatial attention may influence Chinese reading in a “between-character” way. That means the visual-spatial attention deficit in Chinese dyslexia may disturb the visual-spatial attention spotlight sweeping across characters rather than letters, which will decrease the reading accuracy. In conclusion, although Chinese children with DD showed the same performances in the conjunction search task as alphabetic languages children with DD, the mechanism of visual-spatial attention was different. In alphabetic languages, the visual-spatial attention of dyslexic children is related to phonological decoding, and it could influence the reading at the letter level. Conversely, the visual-spatial attention of Chinese dyslexic children is related to visual-orthographic skill, and it could influence the reading at the character level.

### Limitations and future studies

The present study found visual-attention deficit in Chinese dyslexic children. However, whether this deficit was the cause of the Chinese dyslexia is still unclear. Evidence showed that learning to read could enhance the activity of early visual cortex (Dehaene et al., 2010). This finding suggested that the deficit in visual processing in Chinese children with DD might be just the consequence of the lacking reading experience. A reading level control group is needed in future studies to clarify this question. The researchers proposed that the deficit in visual-spatial attention could root in the weakened or abnormal magnocellular input from early visual cortex or LGN to the dorsal visual stream (Stein & Walsh, 1997; Vidyasagar & Pammer, 1999; 2010). Some studies also found that the Chinese children with DD had poor performances in the coherent motion task (Meng, Cheng-Lai, Zeng, Stein, & Zhou, 2011; Qian & Bi, 2014) and higher contrast thresholds in low spatial frequency grating, which was magnocellular sensitive stimuli (Zhao, Qian, Bi, & Coltheart, 2014). One training study showed that the deficit magnocellular-dorsal pathway might be the cause of Chinese children with DD (Qian & Bi, 2015). Thus, more research about whether the magnocellular-dorsal pathway deficit is the cause of developmental dyslexia in Chinese is needed.

In conclusion, the present study found the Chinese children with DD had deficits in visual-spatial attention. The visual processing was correlated with the rapid picture naming, and the rapid orienting of visual-spatial attention was correlated with the component of the orthographic skill. The ability of visual-spatial attention was a special predictor of Chinese reading.

